# Cortical gradients during naturalistic processing are hierarchical and modality-specific

**DOI:** 10.1101/2022.10.15.512379

**Authors:** Ahmad Samara, Jeffrey Eilbott, Daniel S. Margulies, Ting Xu, Tamara Vanderwal

## Abstract

Understanding cortical topographic organization and how it supports complex perceptual and cognitive processes is a fundamental question in neuroscience. Previous work has characterized functional gradients that demonstrate large-scale principles of cortical organization. How these gradients are modulated by rich ecological stimuli remains unknown. Here, we utilize naturalistic stimuli via movie-fMRI to assess macroscale functional organization. We identify principal movie gradients that delineate separate hierarchies anchored in sensorimotor, visual, and auditory/language areas. At the opposite/heteromodal end of these perception-to-cognition axes, we find a more central role for the frontoparietal network along with the default network. Even across different movie stimuli, movie gradients demonstrated good reliability, suggesting that these hierarchies reflect a brain state common across different naturalistic conditions. The relative position of brain areas within movie gradients showed stronger and more numerous correlations with cognitive behavioral scores compared to resting state gradients. Together, these findings provide an ecologically valid representation of the principles underlying cortical organization while the brain is active and engaged in multimodal, dynamic perceptual and cognitive processing.

**Highlights:**

- Movie-fMRI reveals novel, more granular principles of hierarchical cortical organization
- Top movie gradients delineate three separate perception-to-cognition hierarchies
- A distinctive third gradient in movie-watching is anchored by auditory/language regions
- Gradient scores demonstrate good reliability even across different movie stimuli
- Movie gradients yield stronger correlations with behavior relative to resting state gradients

## 1. Introduction

Decades of neuroscience research have studied how local cortical systems integrate simple sensory elements into complex cognitive experiences. Perhaps the most famous example involves the visual system, from the tuning of primary visual cortex neurons to the perception of complex objects (e.g., faces) along a vision-to-perception occipitotemporal visual pathway (Felleman and Van Essen, 1991; Goodale and Milner, 1992; Hubel and Wiesel, 1962; Mishkin and Ungerleider, 1982; Patterson et al., 2007; Visser et al., 2012). Findings like these, and from neuroanatomical and neuroimaging studies, have informed the conceptualization of a macroscale sensory-to-cognition hierarchy of brain organization. For example, Mesulam hypothesized that cognition arises from the integration of information from modality-specific cortical areas (Mesulam, 1998), and a wave of recent studies have investigated these concepts using functional neuroimaging data (Buckner and DiNicola, 2019; Buckner and Krienen, 2013; Deco and Kringelbach, 2017; Margulies et al., 2016).

### 1.1. Functional connectivity gradients

Functional connectivity (FC) refers to the pairwise correlation between brain areas based on fluctuations in BOLD-signal time-courses over time (Biswal et al., 1995). FC is often represented in matrix form in which each column/row represents a connectivity pattern between one region of interest and all other regions. Theoretically, there could be as many unique connectivity patterns (i.e., dimensions) in this matrix as there are regions. Dimensionality reduction techniques assume that the connectivity of different brain regions sits on a lower-dimensional manifold that does not visit the entire higher-dimensional space. Such techniques have long been applied to FC, for example, to identify networks in a data-driven way (Beckmann et al., 2005; Dobromyslin et al., 2012).

Recent studies have introduced diffusion map embedding, a nonlinear dimensionality reduction algorithm, to resting state fMRI data (Langs et al., 2015; Langs et al., 2010; Langs et al., 2011; Margulies et al., 2016). Rather than considering brain networks or regions with distinct boundaries between them, diffusion embedding captures change across the cortical surface based on a certain feature (e.g., similarity in functional connectivity). This method yields a principal gradient that is anchored at one end by unimodal cortical regions and on the other end by transmodal association regions (primarily the default mode network), embodying the sensory-to-cognition conceptualization suggested by Mesulam and others (Buckner and Krienen, 2013; Margulies et al., 2016; Mesulam, 1998). Subsequent work has demonstrated that these FC gradients are conserved across species, similar across analytical methods, and reproducible across datasets (Dong et al., 2021; Hong et al., 2020; Margulies et al., 2016; Shafiei et al., 2020; Xu et al., 2020). They are largely unchanged by sleep deprivation (Cross et al., 2021), but are sensitive to acute pharmacological manipulation (Girn et al., 2022) and at least when focusing on subcortical regions, macroscale cortical gradients show some task versus rest differences (Tian et al., 2020). Overall, despite the fact that this line of research is largely driven by questions about the organizational relationship between sensory and higher-order brain regions, almost all gradient analyses to date have been applied to resting state data, and the effect of active, multimodal neural processing on gradient structure has not been explored.

### 1.2. Gradients and naturalistic imaging

Starting with the assumption that intrinsic FC gradients capture meaningful principles of large-scale brain organization (unimodal-to-transmodal integration across distinct modalities) we wanted to study gradients during active processing that would maximally drive exogenous or evoked functional connectivity patterns. An ideal acquisition state would *i*) engage the brain as a whole rather than targeting a specific brain area or set of areas, *ii*) employ a stimulus or set of stimuli that probe processes along the cognitive hierarchy, ranging from “simple” perception all the way to abstract thinking, social processing, memory and abstract reasoning, *iii*) involve multiple distinct sensory modalities to differentially engage unimodal brain areas, and *iv*) represent real-world content with ecological validity and dynamic context rather than strictly controlled events with sparse temporal order. A naturalistic imaging paradigm such as movie-watching emerges as a useful state that meets these criteria (Hasson et al., 2004; Lahnakoski et al., 2012; Leopold and Park, 2020; Nastase et al., 2020; Redcay and Moraczewski, 2020; Sonkusare et al., 2019; Vanderwal et al., 2019). We hypothesized that functional connectivity gradients derived from movie-fMRI data would reveal new information about cortical brain organization during whole-brain, naturalistic processing and could provide unique tools with enhanced ecological validity.

To test this hypothesis, we first computed cortical FC gradients using movie-watching fMRI data from the Human Connectome Project 7T release (Van Essen et al., 2013). Next, we compared movie-watching gradients to those from resting state data based on topography and test-retest reliability. Finally, we examined brain-behavior correlations and predictive modeling of cognitive, emotional and motor scores in the two conditions. As such, this is the first study to characterize gradient-based principles of functional cortical organization under naturalistic conditions.

## 2. Methods

### 2.1. Data

#### 2.1.1. Participants

All analyses used the HCP 7T release dataset which includes resting state and movie-watching fMRI data from 184 healthy adult participants (122 females, mean age 29.4 ± 3.3). We included subjects who: *i*) had complete functional and behavioral measures of interest), *ii*) passed a two-step motion mitigation, and *iii*) had ≥ 480 volumes (≥ 8 minutes) remaining per functional run. This yielded 95 participants (58 females, mean age 29.5 ± 3.3). The HCP data contain sets of siblings and twins, and these 95 participants represent 64 unique families. All imaging and behavioral data used here were anonymized and made publicly available by the HCP. Participant informed consent forms were approved by the Washington University Institutional Review Board.

#### 2.1.2. Imaging data

Imaging was performed on a 7 Tesla Siemens Magnetom scanner with a Nova32 head coil at the Center for Magnetic Resonance Research at the University of Minnesota. All functional runs were collected using a gradient-echo planar imaging sequence with the following parameters: TR = 1000 ms, TE = 22.2 ms, flip angle = 45 deg, field of view = 208 × 208 mm, matrix = 130 × 130, slice thickness = 1.6 mm; 85 slices; 1.6 mm isotropic voxels. Full details are available at https://www.humanconnectome.org/study/hcp-young-adult/document/1200-subjects-data-release.

#### 2.1.3. Scanning conditions

Four scanning sessions were completed over 2 days and each session started with a resting state acquisition. Sessions 1 (day 1) and 4 (day 2) also included 2 movie-watching acquisitions each (see Supplementary Fig. 1 for schematic). The direction of phase encoding alternated (PA; Rest 1, Rest 3, Movie 2, Movie 3 and AP; Rest 2, Rest 4, Movie 1, Movie 4). Rest runs were each 15:00 (min:sec; 900 TRs). Movie runs varied from 15:01 (901 TRs) to 15:21 (921 TRs).

##### Movies

Participants viewed 4 or 5 different movie clips per movie run. Two types of movies were used: short independent films under Creative Commons licensing (Movie 1 and Movie 3) and clips from Hollywood feature films (Movie 2 had films from 2001-2010, and Movie 4 from 1980-2000). Clip duration varied from 1:04 (64 TRs) to 4:19 (259 TRs) and 20s of rest occurred between clips. All 4 movie runs ended with the same 1:24 (84 TRs) Vimeo clip, and they all started and ended with 20 seconds of rest. Movies were viewed on a 1024 × 768, 4:3 aspect ratio screen rear-projected via a mirror mounted on the top of the head coil, and audio was delivered using Sensimetric earbuds.

##### Rest

Rest fMRI data were acquired with eyes open. A bright cross-hair for fixation was shown on a dark background.

### 2.2. Preprocessing

#### 2.2.1. HCP preprocessing

We used fMRI data that already underwent HCP’s minimal preprocessing pipeline (Glasser et al., 2013). No slice timing correction was performed, spatial preprocessing was applied, and structured artifacts were removed using ICA + FIX (independent component analysis followed by FMRIB’s ICA-based X-noiseifier). Data were represented as a timeseries of grayordinates in CIFTI format (i.e., cortical surface vertices and subcortical standard-space voxels). The first 10 volumes were discarded to allow the magnetization to stabilize to a steady state.

#### 2.2.2. Head motion

To improve cross-condition comparisons, we implemented extra steps to further mitigate the effects of head motion on functional connectivity. Subjects with an overall mean framewise displacement (FD) of > 0.2 mm in any of the 8 functional runs were excluded from all analyses. We then implemented volume censoring (i.e., motion scrubbing) on all rest and movie runs, removing volumes with FD > 0.3 mm, along with 1 volume before and 2 volumes after (Power et al., 2014; Power et al., 2015; Yan et al., 2013).

#### 2.2.3. Cross-condition optimization

All inter-clip rest epochs + 10 subsequent volumes were removed from all movie runs (Supplementary Fig. 1). Full runs were z-scored (as part of the preprocessing), but individual movie segments were not z-scored separately prior to splicing. A potential consequence of this together with motion scrubbing was that some subjects could end up with too few data points to yield reliable FC measures. We thus excluded any subject with < 480 volumes (60%) in any of the 8 main runs following these interventions. The number of volumes for corresponding pairs of runs was then matched within every subject, meaning that Rest runs for some subjects were sometimes truncated. In the few run-pairs where Rest contained fewer volumes than Movie, we retained more rest volumes in the other 3 rest runs instead of cropping the movie runs. This was done to preserve the temporal dynamics in the movie data.

### 2.3. Analyses

#### 2.3.1. Functional connectivity matrices

Concatenated vertex-wise functional timeseries were averaged based on their assignment to the Schaefer-1000 functional parcellation for each subject (Schaefer et al., 2018). For consistency with previous gradient work in resting-state data, we limited our analyses to cortical regions. After applying the parcellation, and as noted previously by Hong et al., (2020), two parcels returned values of not-a-number because no vertices were assigned to those parcels as they were very small (specifically, parcels 533 and 903) (Hong et al., 2020). They were subsequently excluded, leaving 998 parcels that was used in all subsequent analyses. We then correlated all possible pairs of parcel timeseries to build a 998 × 998 FC matrix per condition per subject. Additionally, we computed group-level FC matrices for Rest and Movie by Fisher’s z-transforming and averaging the subject-level FC matrices and transforming the resulting matrices back to correlation coefficients. The group-level FC matrices (based on an average of 2639 volumes, 44 minutes of data per subject) were the basis for the main analysis. To examine the effect of spatial resolution on gradient topography in Movie, we implemented the same gradient pipeline using both the Schaefer-300 and 600 parcellations. For the reliability analyses, we also computed individual subject-level FC matrices based on the test (concatenated Rest 1 and 2, and concatenated Movie 1 and 2) and retest (concatenated Rest 3 and 4, and concatenated Movie 3 and 4) datasets.

#### 2.3.2. Gradient analysis

##### Group-level gradients

Methods closely followed and used code from Margulies et al. (2016) and Vos de Wael et al. (2020) (Margulies et al., 2016; Vos de Wael et al., 2020). The group-averaged Rest and Movie FC matrices were thresholded row-wise at 90% (i.e., the actual correlation values for the above-threshold cells were retained). This process results in sparsity and asymmetry in the thresholded matrices. We next computed a new affinity matrix that captures the inter-area similarity in FC for each condition, using the cosine similarity function. The affinity matrices were then decomposed using diffusion embedding (DE), a non-linear dimensionality reduction algorithm, yielding a separate set of manifolds (i.e., gradients) for rest and movie (Coifman et al., 2005). DE was implemented using the Matlab version of BrainSpace toolbox (Vos de Wael et al., 2020). Compared to other non-linear dimensionality reduction algorithms, the DE algorithm is robust to noise and computationally inexpensive. Notably, the algorithm is controlled by a single parameter α, which controls the influence of density of sampling points on the manifold (α = 0, maximal influence; α = 1, no influence). In this study, we used the default setting of α = 0.5, a choice that retains the global relations between data points in the embedded space (Margulies et al., 2016). The FC gradients are ordered based on the amount of variance explained, and each cortical parcel is assigned a score on each gradient. Since the direction of each gradient is randomly determined, we used a sign flip function to align corresponding rest and movie gradient pairs based only on direction.

##### Group-level PCA components

To compare diffusion embedding to a different dimensionality reduction method, we also applied a basic principal component analysis (PCA) to the same FC matrices thresholded at 90%. The goal of this analysis was simply to explore whether the overall topography of movie gradients would be similar when using a different (and in this case, linear) dimensionality reduction approach. We limited cross-method statistical comparison to computing the Pearson’s correlation coefficient between DE and PCA gradients at each gradient for the first 10 gradients. This was the only analysis in which PCA components were used.

##### Individual subject-level gradients

To provide a level of quality control after the BrainSpace toolbox pipeline, we plotted subject by parcel scores of the top gradients, creating a 95 × 998 matrix for each gradient for visual inspection. A small number of rows (subjects) had markedly low variability in gradient scores across parcels in some gradients. Gradient scores of top gradients for those subjects were projected on the cortical surface and showed unusual gradients that were anchored by one or a few parcels that had unusually high scores. Additionally, other individual-level gradients demonstrated similar organization to group-level gradients but were hard to visualize due to one or a few parcels with outlier scores that artificially inflated the range of these gradients. To handle these outlier gradients and scores on analyses conducted at the individual-subject level, a data-driven two-step outlier removal process was implemented for each subject. This process was limited to the first 10 gradients. First, whole gradients were considered outlier gradients and removed if their maximum absolute gradient score divided by the median score for that gradient was > 3 standard deviations above the median score of the corresponding group-level gradient. This step resulted in the removal of 64 out of 950 gradients (95 subjects × 10 gradients) from 39 different subjects. Second, (absolute) gradient scores > 3 standard deviations above the median score for that subject’s gradient were identified as outlier scores and set to 0. Individual-level gradients were then aligned to group-level gradient templates from the same condition using Procrustes alignment without scaling.

#### 2.3.3. Decoding the organization of top Movie gradients

We used the NeuroSynth database to examine neuroscientific terms associated with regions of interest created from 10-percentile bins at the unimodal end of the first 3 group-level gradients in Movie. Considering the arbitrary polarity of gradients, we manually selected the pole opposite from the hetermodal frontoparietal/default pole for each gradient to create these bins. Word clouds were used to represent the term weights for each gradient.

#### 2.3.4. Test-retest reliability

To maximize the amount of data for reliability estimates across scan days, we computed gradients at the individual subject-level for a subset of subjects who had ≥ 600 volumes (10 minutes) per run for all 8 runs (following motion mitigation and cross-condition optimization steps, n=67, 40 females, mean age = 29.6 ± 3.3, 49 families). For each condition for each subject, data were divided into test and retest datasets, each consisting of 2 concatenated runs acquired on the same day: Rest test [Rest 1 + 2], Rest retest [Rest 3 + 4], Movie test [Movie 1 + 2], and Movie retest [Movie 3 + 4]. We computed individual subject-level gradients based on each of the four datasets. Since the manifold spaces in Rest and Movie were different, each set of individual-level gradients was aligned to the group-averaged gradients of the same condition using Procrustes alignment. Intraclass correlation coefficient (ICC), defined as the between-subject variability divided by the sum of within- and between-subject variability was computed using ICC model 2, denoted as ICC(2,1) (Noble et al., 2021; Shrout and Fleiss, 1979). Gradient reliability as a function of data amount was evaluated by varying the amount of data used to compute the individual-level gradients (1-20 minutes, 1-minute increments). ICC at each time point was computed, and two-tailed t-tests were performed comparing cross-condition ICC reliability at 10 and 20-minute marks. To test the hypothesis that movie and rest gradients are distinct, we also computed ICC values for scans across conditions (Rest to Movie, and vice versa). One-way ANOVA test was performed on within- and cross-condition ICC values.

#### 2.3.5. Correlation between gradient scores and behavioral measures

A parcel’s score along a given gradient indicates the relative position of that parcel along that gradient based on its connectivity pattern, or put differently, is akin to that parcel’s weighting within that organizational component. At the individual subject level, the gradient score of a parcel might contain idiosyncratic information that could be linked to phenotypic traits (Cross et al., 2021; Hong et al., 2020). To test for cross-condition differences in gradient-behavior relationships, we performed cortex-wide univariate correlations between gradient scores in Rest and Movie and behavioral scores.

##### Behavioral data

Twelve cognition, 19 emotion and 4 motor scores were used (see supplementary Table 1 for full list) from the HCP data. To deal with potential intercorrelations of scores within each domain, we performed PCA on z-scores of variables within each domain to yield one latent variable per domain (Finn and Bandettini, 2021; Vanderwal et al., 2021). PC1s explained 26%, 30%, and 39% of the variance in the cognition, emotion, and motor data, respectively, and each participant’s component score for each PC was used as the behavioral measure of interest.

##### Gradient score-behavior correlations

For each of the top 4 gradients, we computed the Pearson correlation coefficient between 2994 (998 parcels × 3 behavioral variables) pairs of individual subject-level gradient scores and behavioral scores across the n = 95 subjects in Rest and Movie separately. Differences in brain-behavior correlations for the first 4 gradients between Rest and Movie were assessed using t-tests of absolute r-values across the various pairs of conditions. Absolute r-values here account for both positive and negative correlations between gradient and behavioral scores. Additionally, the number and topography of parcels with |r| > 0.2 (p < 0.05) with each behavioral measure were examined and compared across conditions. This threshold was implemented for visualization, and we did not correct for multiple comparisons in this instance.

##### Brain-behavior predictions

To investigate whether the gradient organization during movie-watching relates to behavior at the individual subject level, we compared the ability of movie gradients and rest gradients to predict behavior. Both single and combined gradient maps were used as predictors. We directly fitted a ridge regression model (R glmnet package, https://glmnet.stanford.edu/) with training subjects using full gradient maps for each input-output pairing and applied the model to testing subjects in a 10-fold cross-validation framework. Default parameters were used (i.e., lambda = NULL and alignment = “lambda”). Familial relatedness was accounted for by confining siblings to either the training or the testing folds. One hundred values of the λ parameter (degree of penalization) in the ridge regression were automatically chosen by the glmnet algorithm and tested at each fold. The λ value that minimizes the mean squared error of cross validation was chosen for that model for that fold. The fits are then aligned using lambda values derived using the full data set, but applied across all the fold-level models, and a single best-performing “harmonized” lambda value is selected. We performed 100 iterations of this process for each model to assess if its accuracy was sensitive to different folds. Prediction accuracy was evaluated by calculating the Pearson’s correlation coefficient between the predicted and true scores, as well as mean absolute error (MAE) for each model iteration. To assess the statistical significance of prediction accuracies, we first generated a null distribution of expected accuracies due to chance by shuffling behavior scores with respect to gradient maps and reperforming the predictive modeling pipeline 10,000 times for each input-output pairing. We then calculated a p-value according to permutation testing for each model.

The primary focus in the predictive modeling analyses was on movie gradients vs. rest gradients as above. A previous study had shown that movie FC data outperforms rest FC data for predictive modeling (Finn and Bandettini, 2021), while a separate study showed that rest gradient scores outperform rest FC scores (Hong et al., 2020). It thus seemed likely that movie gradients would outperform movie FC, but we at least wanted to assess that here. Consequently, we also included full rest and movie FC matrices as inputs in our predictive modelling, noting the significant difference in size between these inputs, as FC matrices are 996,004 (998 × 998) and gradient maps are only 998 or 3,992 (998 × 4) in the case of combined gradients.

Paired t-tests were used to compare the r and MAE values from the 100 iterations between Rest and Movie for each gradient map/connectome input that yielded predictions that differed significantly from the null distribution. False discovery rate was used to correct for multiple t-tests used to compare predictions within each behavioral domain.

### 2.4. Data and code availability

MRI and behavioral data are publicly available via the HCP. All code used to perform the analyses in this study is available at https://github.com/tvanderwal/naturalistic_gradients_2022.

## 3. Results

Group-averaged FC matrices contained an average of 2,639 volumes (44 min) of data per subject, and Rest and Movie data amounts were matched within subjects. Using these “optimized” FC matrices, diffusion embedding was applied to extract group-level FC gradients (n = 95, Fig. 1). The first 10 gradients explained similar variance in Rest and Movie (t-test: t = 0.9677, p = 0.3334; Fig. 1A). The first 3 group-level gradients explained 11.53%, 9.84% and 8.14% variance in Rest, and 10.45%, 9.76% and 8.03% variance in Movie. Next, we examined the similarity between the first 10 gradients across conditions using Pearson correlation coefficient (Fig. 1B). Gradients 1 and 2 showed the highest similarity between Rest and Movie (r = 0.87 and 0.86, respectively), rest-G3 was strongly correlated with movie-G4 (r = 73), and all other correlations were fairly low (r < 0.67). The strongest correlation between movie-G3 and any of the rest gradients was r = 0.46.

**Fig. 1.**
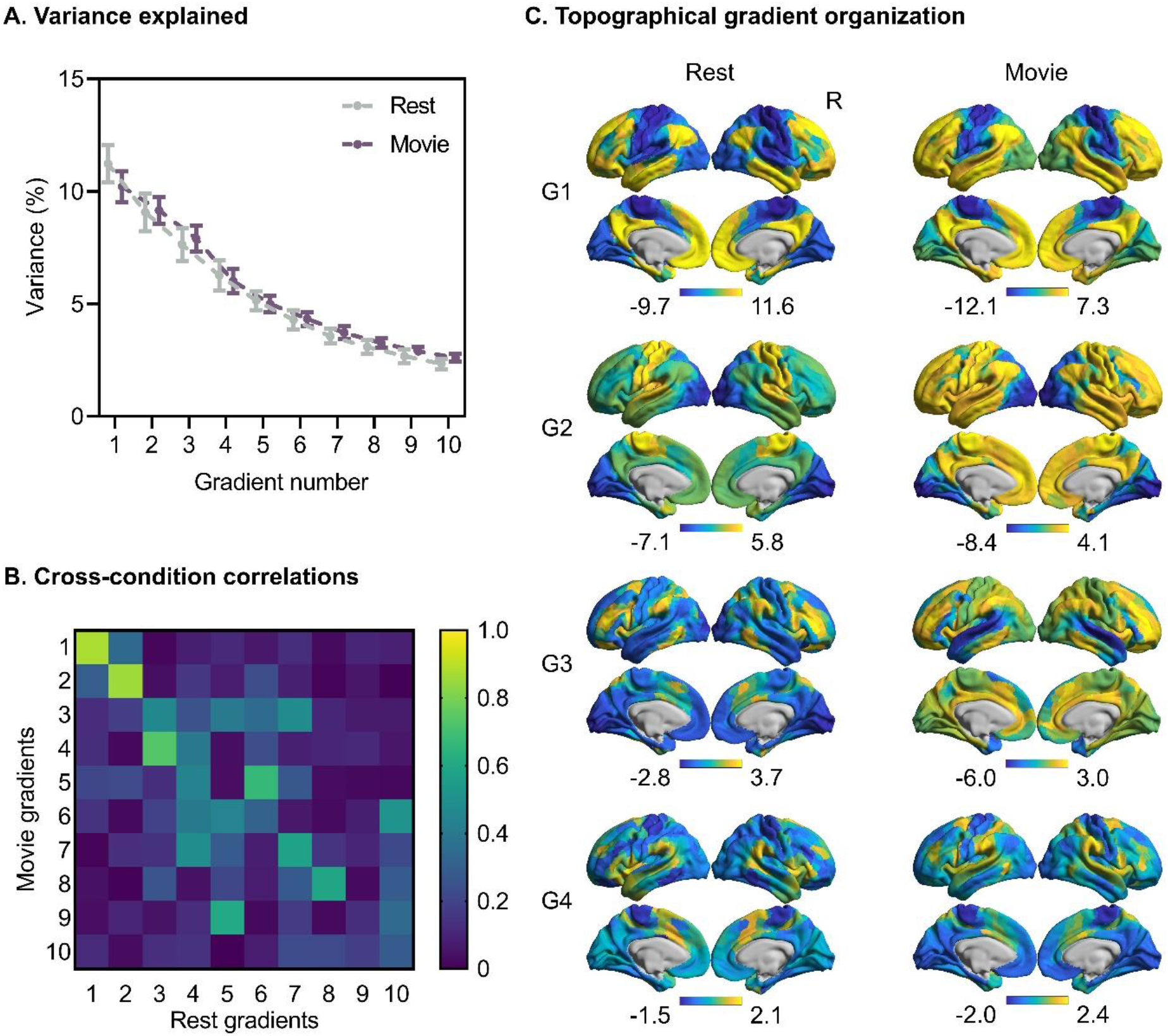
Topography of the top four rest and movie FC gradients (N = 95). **(A)** Scree plot showing variance explained by the first 10 gradients, which was similar between Rest and Movie (t-test: t = 0.9677, p = 0.3334). Error bars indicate 1 standard deviation. **(B)** Pearson correlations between the first 10 gradients in Rest and Movie. Only the first 2 gradients show strong correlations across conditions, movie-G4 is correlated with rest-G3, and notably, movie-G3 does not look similar to any rest gradient. **(C)** Gradient scores projected on the cortical surface show similarities and differences in topography between Rest and Movie. Rest-G1-4 replicate previous work. In movie-G1, visual regions are situated midway between unimodal and heteromodal regions, and auditory regions coalesce with default mode and frontoparietal networks. For movie-G2, most transmodal and nonvisual primary regions have similar scores, creating a visual-nonvisual axis. Movie-G3 is anchored by auditory and language areas in the superior temporal gyrus and sulcus (including Wernicke’s area), Broca’s area, posterior middle frontal gyrus, and area 55b. Similar to rest-G3, movie-G4 is anchored by task-positive areas of the dorsal and ventral attention networks. Gradients are shown here each in its own scale to better highlight topography, but Supplementary Fig. 2 shows gradients in the same scale to better represent the relative variance.

### 3.1. Gradient Topography

When projected onto the cortex, gradient scores reveal both topographical differences and similarities between Rest and Movie (Fig.1C; gradients shown using the same scale are shown in Supplementary Fig. 2). Rest gradients replicate those previously described (Hong et al., 2020; Huntenburg et al., 2018; Margulies et al., 2016): rest-G1 forms a unimodal-to-transmodal hierarchical axis with visual, sensorimotor and auditory regions together, anchoring this axis’s unimodal pole. Rest-G2 separates regions based on modality, from visual to sensorimotor/auditory regions. In Movie, G1 also captures a unimodal-to-transmodal axis. However, only sensorimotor regions anchor the unimodal pole of this gradient, and visual regions are shifted to the middle of movie-G1. Also, in movie-G1, auditory regions are part of the heteromodal end of the gradient together with default network regions. Movie-G2 captures a second unimodal-to-transmodal axis in which visual regions are uniquely isolated, creating a visual-nonvisual gradient. Auditory regions form the basis of a unique axis captured by movie-G3. This unique movie gradient is anchored by auditory and language areas in the superior temporal gyrus and sulcus (including Wernicke’s area), Broca’s area, posterior middle frontal gyrus, and area 55b. The first 3 movie gradients are each anchored at the transmodal pole by areas that participate in the default mode, limbic and frontoparietal function networks. Rest-G3 and movie-G4 have similar topography running from unimodal and task-negative areas to task-positive attention areas. Together, the top 3 movie gradients are modality-specific, hierarchical gradients. The organization of top movie gradients remains largely unchanged when using coarser parcellation resolutions (Supplementary Fig. 3).

#### PCA vs Diffusion Embedding (DE)

Movie gradients are highly similar when using PCA (Supplementary Fig. 4). The first 3 PCA components explained, in order, 16.35%, 12.31% and 9.40% variance in Rest, and 15.02%, 10.51% and 7.79% variance in Movie. Correlations of gradients 1-4 between DE and PCA are significantly higher in Movie relative to Rest, and the difference remains significant when the first 10 gradients are tested (Supplementary Fig. 4).

### 3.2. Organization of sensory regions within rest and movie gradients

Gradient scores within networks are plotted for the top 4 gradients in Rest and Movie (Fig. 2). The hierarchical nature of movie G1-3 is evident when comparing mean scores across networks (Fig. 2A). Gradient scores for G1-3 were also plotted to create 3-dimensional gradient space for Rest and Movie (Fig. 2B). Parcels were color-coded based on their assignment to functional networks. In Rest, only G1 is hierarchically organized, and sensorimotor and auditory parcels from the somatomotor network are grouped together along all gradients. In Movie, the auditory parcels separate out from the somatomotor network, and G1-3 converge towards the DMN and frontoparietal. The parcels that move the greatest distance in gradient space from Rest to Movie are located along the superior temporal sulcus (STS) (Fig. 2D). See Github repository for animation visualizing the amount of modulation between states (https://raw.githubusercontent.com/tvanderwal/naturalistic_gradients_2022/main/resources/naturalistic_gradients.gif). To examine the functional hierarchy along movie gradients in reference to the hierarchical organization of rest-G1, we examined the rank order of parcels within movie G1-4 against their rank along rest-G1 (Fig. 2C). Movie G1-3 but not G4 resemble rest-G1 based on parcel rank (most parcels follow the y = x line trajectory). All but one modality move up the hierarchy (i.e., falls below the y = x line) in G1-3, demonstrating modality-specific organization. Functional segregation along sensory modalities is evident such that Movie G1 is associated with “pure” sensorimotor functions, movie-G2 with visual functions, and movie-G3 with a complex set of language and auditory functions. Term-based decoding of these regions using Neurosynth reflects clear functional and regional specification of each movie gradient (Fig. 3).

**Fig. 2.**
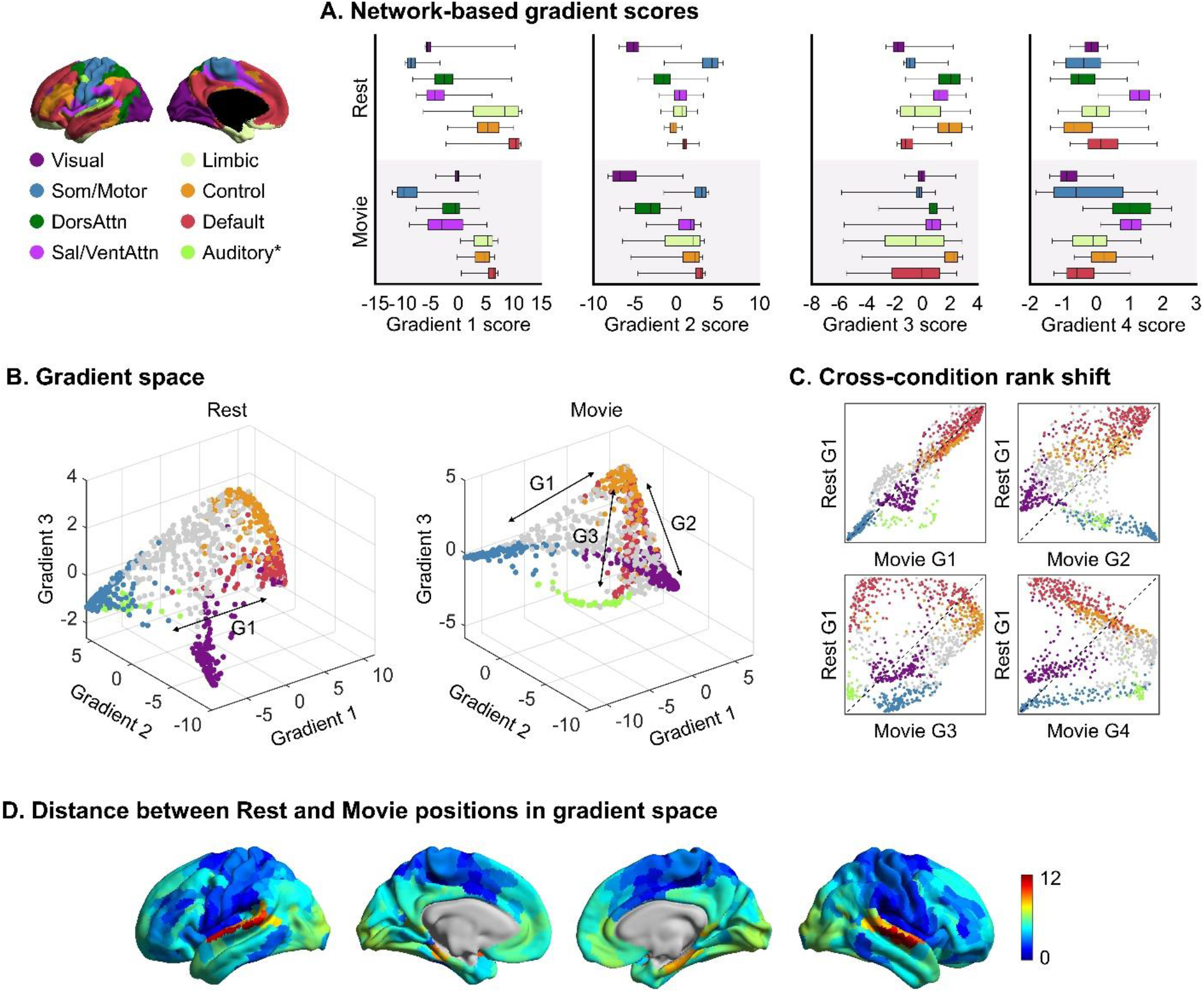
Movie gradients are modality specific. Parcel-based gradient data were summarized based on the 7 canonical networks (Yeo et al., 2011). An auditory network (neon green) was defined based on movie-G3 scores, and is shown in B. **(A)** Boxplots of gradient scores within the 7 networks show where each network falls along the gradients. **(B)** Scatterplots of the top 3 gradients (i.e., gradient space) in Rest and Movie color-coded by parcel assignments to FC networks from A. In Rest, somatomotor and auditory networks behaved as one unit, whereas in Movie, the two networks decoupled along gradients 1 and 3. Rest-G1 runs from auditory, sensorimotor, and visual regions to heteromodal regions, replicating the “principal gradient” of unimodal-to-heteromodal functional hierarchy. In Movie, all 3 gradients represent hierarchical axes. The heteromodal end of movie-G1, G2 and G3 is composed of both default and frontoparietal networks, and each gradient is anchored in one modality at the unimodal end: sensorimotor in movie-G1, visual in movie-G2, and auditory in movie-G3. **(C)** Scatterplots showing the rank of each parcel along movie-G1-4 relative to their rank along rest-G1 (i.e., rest functional hierarchy). The triangles below and above the dotted line (y = x line) represent moving up and down the hierarchy in Movie, respectively. The distribution of parcels along the y = x line in movie-G1-3 (but not movie-G4) recapitulates the hierarchical nature of these gradients. **(D)** Distance between Rest and Movie parcel positions in 3D gradient space is projected onto the cortical surface. Colorbar represents Euclidean distance units. Regions from the superior temporal sulcus show markedly higher distance than other regions, followed by visual and default regions. An animation showing the modulation of brain regions in gradient space from Rest to Movie can be viewed at https://raw.githubusercontent.com/tvanderwal/naturalistic_gradients_2022/main/resources/naturalistic_gradients.gif.

**Fig. 3.**
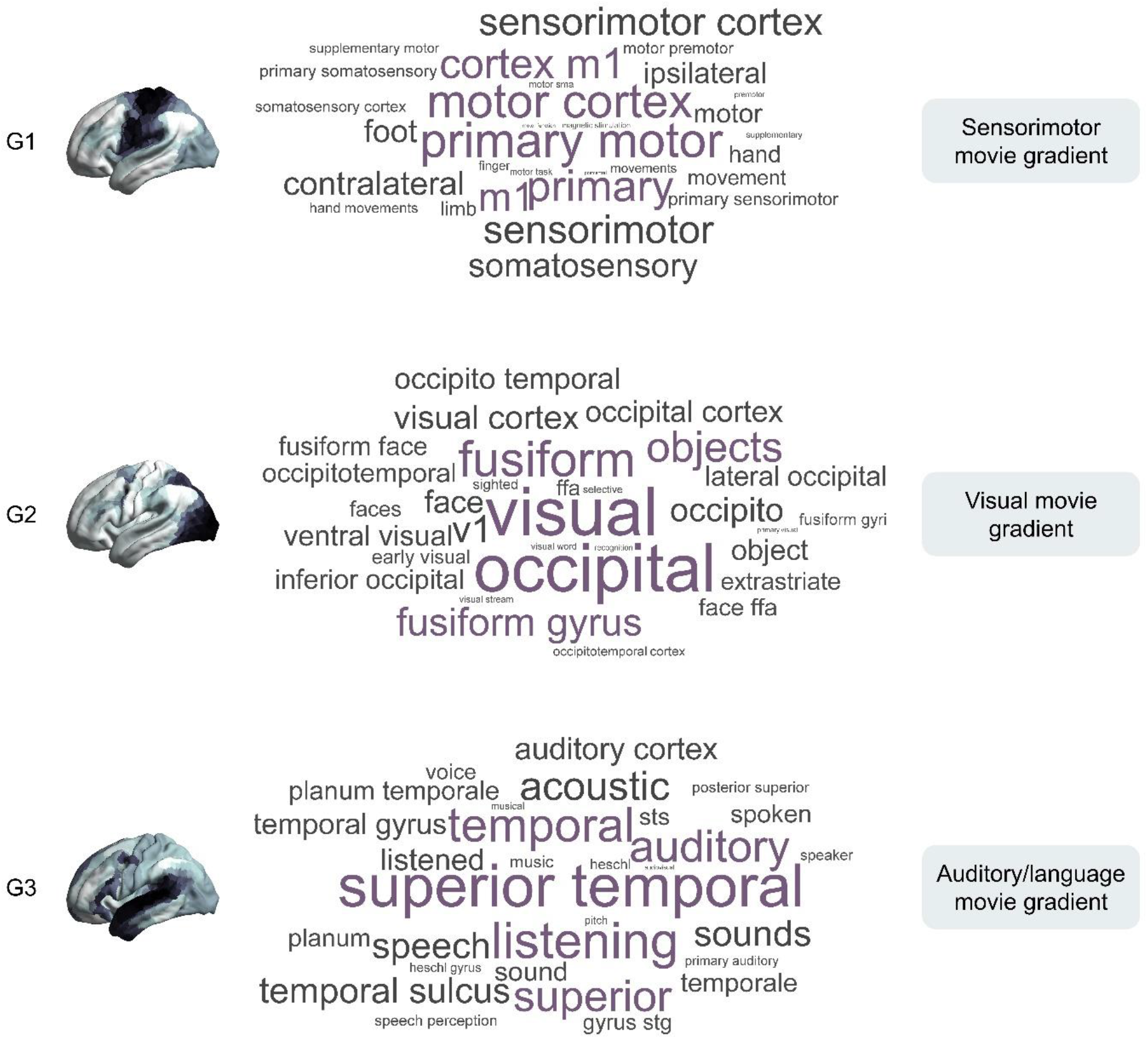
Meta-analytic decoding of movie gradients. Outputs from Neurosynth term-based decoding of the lowest 10% of gradient scores per gradient underscore functional and regional differentiation by modality for each of the top 3 movie gradients.

### 3.3. Test-retest reliability of movie and rest gradients

We compared the test-retest reliability of rest and movie gradients using ICC in a subsample of n = 67 with ≥ 600 volumes (10 minutes) per run for all runs (Fig. 4). Both rest and movie gradients had strong TRT reliability. Despite using different movie clips across scans, the reliability of movie gradients 1-4 was not different from Rest at 10 minutes of data (mean ICC in Rest = 0.73; mean ICC in Movie = 0.74; t-test: t = 0.3639, p = 0.7171). Rest gradients reach a higher ICC compared to Movie gradients at 15 min (Rest ICC = 0.83; Movie ICC = 0.81; t-test: t = 2.077, p = 0.0417) The difference between Rest and Movie becomes nonsignificant again at the 20-minute mark (Rest ICC = 0.84; Movie ICC = 0.87; t-test: t = 1.978, p = 0.0521). When mapped onto the cortex, ICC scores for Movie appeared lowest in occipital, temporal, and default network regions (Fig. 4C). Rest and movie gradient reliability were both significantly higher than cross-condition reliability in either direction (p-values < 0.0001) (Supplementary Fig. 5).

**Fig. 4.**
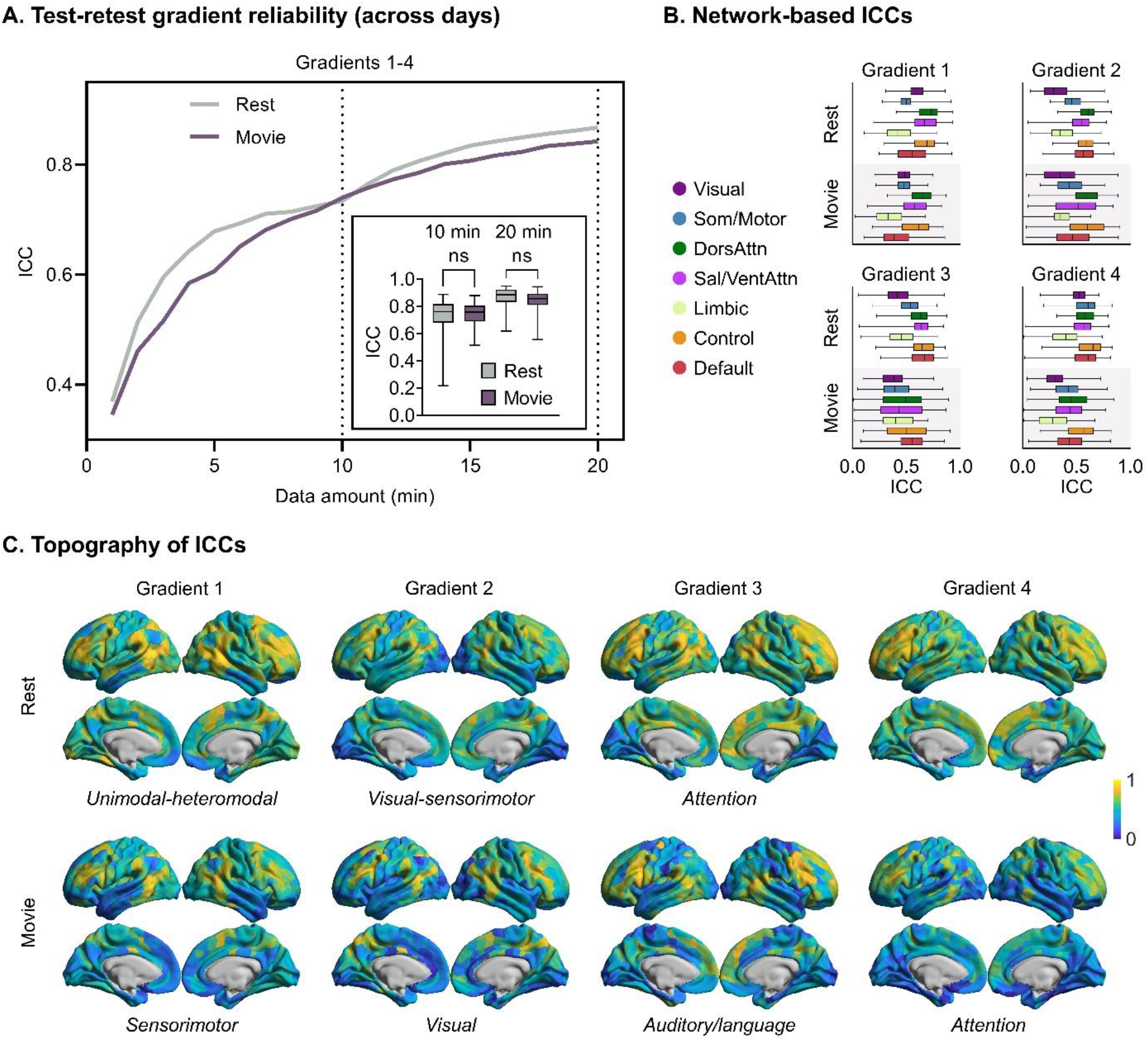
Test-retest reliability of movie and rest FC gradients. **(A)** ICCs for G1-4 (averaged across subjects and gradients) increase as a function of data amount with a similar slope in both Rest and Movie. ICC reaches > 0.6 in Rest and Movie when using 4 and 5 min of data (240 and 300 TRs), respectively. ICC of > 0.8 is achieved with 13 and 14 min of Rest and Movie data, respectively (780 and 840 TRs). Inset shows cross-condition statistical comparisons of ICCs at two arbitrary time-points (10 and 20 min. of data). No significant differences between Rest and Movie are observed. **(B)** Boxplots show ICCs within the canonical networks across the top 4 gradients in each condition. **(C)** ICCs mapped onto the cortical surface show that ICC of gradient scores vary across the brain. For G1 and G2, regions with stronger correlations appear similar across Rest and Movie, and in general, the strongest ICCs appear to include the frontoparietal network and other patchy regions. Ns = P ≥ 0.05.

### 3.4. Brain-behavior correlations

We computed the Pearson correlation coefficient between scores from the first 4 gradient maps with composite (PCA-based) behavioral scores from cognition, emotion and motor domains at each parcel (n = 95). At the parcel level, gradient-behavior correlations for both Rest and Movie were low (maximum r = 0.46 for Movie, r = 0.37 for Rest) (Fig. 5A). In almost all cases, correlations between absolute r-values were significantly stronger for Movie relative to Rest (see Supplementary Table 2). Maps of the correlations showed more regions and stronger correlations for Movie, with regions appearing to cluster by network membership (i.e., not random, especially in Movie) (Fig. 5B and unthresholded maps shown in Supplementary Fig. 6). Additionally, we counted the number of parcels with correlation strength of |r| > 0.2 (P < 0.05 at n = 95). Movie had significantly more parcels above this threshold (Supplementary Fig. 6). When examining brain-behavior correlations within networks, correlations were strongest within the frontoparietal and default networks in movie-G1&2 and visual and dorsal attention networks in movie-G3&4 (Fig. 5C).

**Fig. 5.**
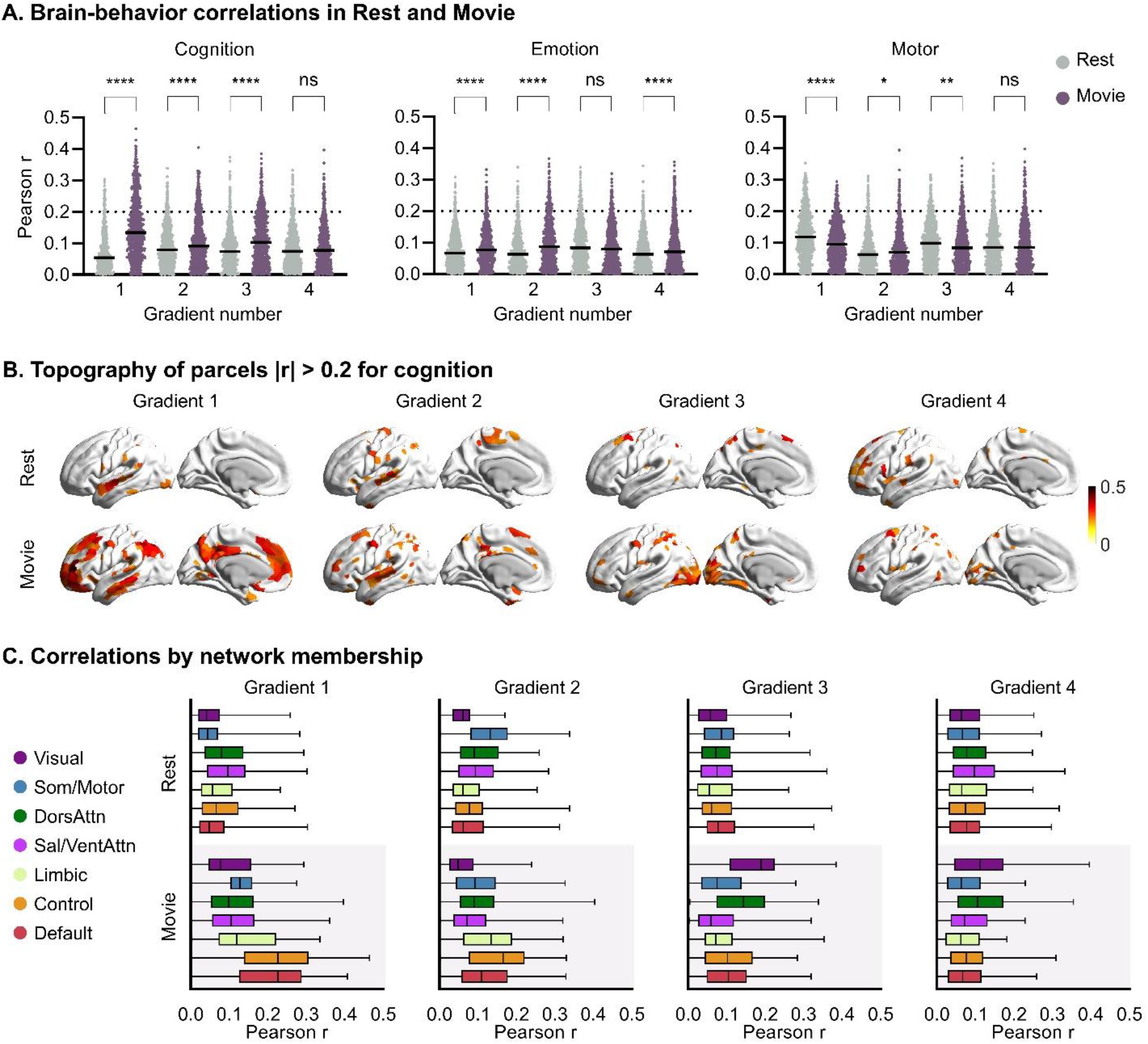
Relating gradient scores to behavioral scores in Rest and Movie, n = 95. **(A)** Cognitive, emotion and motor composite (PCA-based) behavioral scores were used. Absolute values of Pearson’s correlation coefficients are shown for each parcel. Hard line shows mean for each gradient, and r = 0.2 is shown with a dotted line to indicate p < 0.05. **(B)** Correlations (absolute) > 0.2 were projected on the cortical surface. **(C)** Boxplots of Pearson’s correlation coefficients within the 7 canonical networks (Yeo 7-network parcellation) show overall stronger correlations in Movie, especially within default and frontoparietal networks in movie-G1&2 and visual and dorsal attention network in movie-G3&4. ns = P ≥ 0.05; * = P < 0.05; ** = P < 0.01; **** = P < 0.0001.

### 3.5. Gradient-based prediction of behavior in Rest and Movie

To compare brain-behavior predictions at the whole brain, individual subject level across Rest and Movie, we used ridge regression to predict the same composite cognition, emotion and motor scores using single gradient maps, combined gradient maps, and full connectomes (Supplementary Fig. 7). When comparing the Pearson correlation coefficient between predicted and observed scores, significant predictions were found when using movie-G1 and combined movie-G1-4 for cognition, movie-G3 for emotion, and rest connectome for motor. When using mean absolute error to assess model accuracies, only the sensorimotor movie gradient and all movie gradients combined for cognition, and rest FC for motor, remained significantly different from the null models. Within this framework, significant cross-condition differences existed such that Movie outperformed Rest for all gradient-based models that differed significantly from null. Models in some cases yielded negative r-values, indicating overfitting.

## 4. Discussion

An emerging body of literature is using dimensionality reduction techniques with intrinsic functional connectivity data to reveal a gradient space that spans from unimodal to heteromodal cortical regions (Bernhardt et al., 2022; Hong et al., 2020; Huntenburg et al., 2018; Margulies et al., 2016). Here, we build on this work by investigating state-based differences in FC gradients. Specifically, we compare gradients from resting state and movie-watching conditions, the latter providing a brain state during which both sensory and higher-order cognitive processes are dynamically engaged. Movie gradients underscored the hierarchical organization of the cortex observed during Rest but also captured modality-specific granularity (movie G1 = sensorimotor, movie G2 = visual, movie G3 = language/auditory). Even though different movies were used across scans, the test-retest reliability of movie and rest gradients showed no significant difference. Gradient scores–which signify the relative position of brain areas along each gradient– showed stronger correlations with phenotypic traits across subjects in Movie compared to Rest. Here, we attempt to summarize and synthesize notable observations about the movie gradients, what these findings might tell us about brain organization, and how movie gradients could be used in future research.

### 4.1. Default and frontoparietal networks form the heteromodal peak of top three movie gradients

The principal gradient of resting state FC is anchored by unimodal regions on one end and mainly default network regions on the other. Indeed, one of the initial findings of interest was that principal gradient scores were related to the geodesic distance from default network hubs to primary sensory regions (Margulies et al., 2016). In movies, the heteromodal pole of the top three gradients are different insofar as they converge on regions of both the frontoparietal and default networks. That the default network is “isolated” in the resting state principal gradient is not surprising given what we know about unique default network functions in the absence of a directed task (e.g., mind wandering, self-referential processing, future/past thinking, internal representations, etc.) (Andrews-Hanna, 2012; Buckner and DiNicola, 2019; Smallwood et al., 2021). But what does it mean that both frontoparietal and default networks have similar gradient scores across the top gradients during movie-watching?

Recent work using movie fMRI has shown that frontoparietal network function is different during movie-watching than during task, and specifically, that it does not uniquely function as a “switching hub” during movies (Caldinelli and Cusack, 2022; Cole et al., 2013). This is despite the fact that brain states are more numerous and that switching between states occurs more frequently than during rest, likely as a function of the movies themselves (van der Meer et al., 2020). The high test-retest reliability of movie gradients in the frontoparietal network across different movie types (Fig. 4B) suggests that, on average across a compilation of clips, the functional profile of the frontoparietal network is readily generalizable to different movie stimuli.

The role and dynamics of the default network during movie-watching are also still being elucidated, but it has long been implicated in social cognition and narrative processing (Nguyen et al., 2019; Simony et al., 2016; Yeshurun et al., 2021) and overall, it exhibits high levels of FC (Vanderwal et al., 2015). In work emphasizing the pervasive role of memory at all levels of cognitive processing, Hasson et al. (2015) showed that both default and frontoparietal networks contain regions that are near the top of a processing hierarchy based on timescales of information accumulation, and that these regions share relatively long temporal receptive windows (Hasson et al., 2015). In sum, previous work on task-evoked FC changes generally shows that both frontoparietal and default networks are highly engaged during movie watching, indicating that “default” functions such as self-referential processing and social reasoning may occur in concert with executive control functions, and that this interplay or coalescence might emerge specifically in naturalistic contexts that include evolving narratives. The finding that they have similar gradient scores and that they jointly form the heteromodal pole of movie gradients essentially redefines the nature of the hierarchical gradient during naturalistic processing, and raises questions about the extent to which the two networks work together or remain segregated (Fair et al., 2007; Wang et al., 2021). One interesting conceptual question that is sparked by this finding is whether frontoparietal and default networks would share the heteromodal hub in a movie-watching macaque study, or if this intercalated arrangement is an evolutionary product specific to humans.

Taking a network perspective to guide movie gradient interpretation has its limitations, especially considering that the canonical functional networks have been defined in rest (Yeo et al., 2011). The range of gradient scores within each network is wider in Movie compared to Rest suggesting that areas within the same network diverge in their FC patterns during movie-watching. This is most evident in the somatomotor network where movie-watching appears to evoke distinct FC patterns within the sensorimotor and auditory regions to the point of forming separate gradients driven by the two subsystems (gradients 1 and 3, respectively), and similar divergence could well be happening in other networks. Future work using movie-derived networks mapped onto movie gradient space might help make further sense of the cortical gradients, and though here we have tracked network positions from the well-defined resting state to movie, it might be useful to understand how movie-derived networks redistribute in the opposite direction (i.e., from movie gradient space to rest gradient space).

### 4.2. Sensorimotor movie gradient

The overall topography of the first gradient is highly similar between Rest and Movie with some important differences (Fig. 1B&C). While the sensorimotor cortex remains anchored at the unimodal end in Movie, visual and auditory cortices move up the hierarchy (Fig. 2C) rendering movie-G1 a pure sensorimotor-to-heteromodal axis. This trend of primary regions moving up or becoming more “heteromodal” during movies is observed across the gradients overall. The amount that each sensory modality has moved within the sensorimotor movie gradient (and the order of movie-G1-G3) recapitulates the evolutionary and developmental hierarchy of the brain (Dong et al., 2021; Fair et al., 2008; Gilmore et al., 2018; Sydnor et al., 2021; Xu et al., 2020). This adds a “movie gradient” contribution to previous work that has considered the way that gradients relate to both anatomical and evolutionary hierarchies (Sydnor et al., 2021). The sensorimotor focus of movie-G1 is also evident in its term-based meta-analysis (Fig. 3). The differences in gradient 1 organization between Rest and Movie become more significant when examining the correlations between gradient scores and behavioral scores. This sensorimotor movie gradient shows the strongest, most numerous correlations with cognition, with the highest correlations in heteromodal default and frontoparietal networks (Fig. 5B&C). Moreover, this movie gradient outperformed the rest principal gradient for predicting cognitive and emotion scores in a ridge regression cross-validation framework. Given these results, and in light of the growing interest in FC gradients as a framework for biomarker discovery in psychiatry, the sensorimotor movie gradient could provide a useful alternative to the principal gradient in rest for interrogating the biological underpinnings of psychiatric disorders (Hong et al., 2019; Hong et al., 2020; Huntenburg et al., 2018; Xia et al., 2022). Further testing of this idea would require larger datasets that use movie-watching as an acquisition state.

### 4.3. Visual movie gradient

In contrast to rest-G2, which differentiates between cortical areas based on perceptual modality (visual-to-sensorimotor), the second movie gradient is anchored by the visual cortex with almost all other cortical areas forming a non-visual pole. To date, the visual cortex has not been the focus of human gradient work, possibly because it does not stand out in rest gradients, whereas in animal gradient studies, it does. For example, in mice, the visual cortex drives a number of gradients, one of them (murine gradient 3) is a nonhierarchical somatomotor-to-audiovisual gradient that has been shown to have strong correspondence with gene expression (Huntenburg et al., 2021). Joint embedding studies on monkey and human data found that cross-species FC homology recapitulates previously established visual processing order with increasing eccentricity (Xu et al., 2020). The same ordering can be seen here along the lower gradient scores of the visual movie gradient: a local visual hierarchy is reflected in gradient scores as one progresses out from primary visual regions to associative visual regions, providing another example of a hierarchical gradient that follows the topology of the geodesic surface. As such, this visual movie gradient might provide a way to query cortical organization in relation to the visual cortex in humans in a new way.

### 4.4. Auditory and language movie gradient

Movie gradient 3 shows a unique topographic organization not seen in resting state gradients. This gradient is anchored by the full catalog of auditory and language areas, including the superior temporal gyrus and sulcus, posterior middle frontal gyrus, inferior frontal gyrus, area 55b, and Brodmann’s Areas 44 and 45. This complex set of regions is highly similar to an “audiovisual social perception network” shown previously in task-based approaches to movie-watching data (Lahnakoski et al., 2012), during story listening (Huth et al., 2016) and they closely resemble one of the meta-analytic groupings (MAG 2) from a naturalistic meta-analysis (Bottenhorn et al., 2019). It is particularly interesting that the brain regions evoked by multiple social and language-related tasks—and which are connected via different dorsal and ventral language-related white matter tracts (Blazquez Freches et al., 2020; Skeide et al., 2016)—all have similar gradient scores (i.e., movie-G3 goes beyond a “simple” sensory modality). The pole also includes regions in the anterior temporal lobe (ATL). Tract tracing studies have shown convergence of multiple sensory modalities in the ATL (Ralph et al., 2017), and it has been hypothesized to serve as an integration hub for semantic information (Jefferies, 2013). This movie gradient thus uniquely includes both primary and association cortical regions at its sensory end, making its pole more complex than the other movie gradient poles. This auditory/language movie gradient was also the only gradient that showed potential for predicting emotional scores (Fig S5). The heteromodal pole of this gradient is also unique among the movie gradients insofar as it is anchored more by frontoparietal than default network, suggesting that the integration between auditory and frontoparietal regions during movie-watching is distinct from rest. This gradient could provide a unique, comprehensive tool with which to assess functional brain organization as it relates to language and social processing regions. As noted above, the language regions are known to develop later than sensorimotor and visual regions, and it would be interesting to track developmental trajectories of this auditory/language gradient in a longitudinal sample.

We also note that gradient scores in temporal regions seem important across all movie gradients. In the first movie gradient, temporal regions are not grouped with the other sensory regions as they are during rest. Rather, they are grouped together with heteromodal regions, suggesting that during active processing, the connectivity patterns in auditory regions are more similar to heteromodal connectivity patterns. Relatedly, a discrete cluster of regions along the STS are also observed to move the maximal distance within gradient space from Movie to Rest (Fig S3). This distance or cross-state change occurs mainly in the auditory/language movie gradient. Temporal regions are also where we observed the strongest brain-behavior correlations from within the visual movie gradient where they have intermediate gradient scores. These observations suggest that the position of auditory and language regions relative to the rest of the cortex during complex, dynamic neural processing is pivotal, and examining gradient scores in these regions in developmental and psychiatric disorders such as autism, schizophrenia and dyslexia (among others) would be of interest.

### 4.5. Attention gradient

Movie-G4 appears topographically similar to and correlated with rest-G3: both gradients highlight task positive attention networks. The distribution of gradient scores within networks in Fig. 2A shows that in Movie, the task positive/task negative demarcation is even more pronounced. Additionally, movie-G4 correlations with cognition appear to be in the same regions as rest-G3 to cognition. Essentially, we suggest that movie-G4 provides a better representation of the same principle captured in rest-G3 and that given the nature of movie-watching as a robust, dynamic task-state, it is intuitive that a gradient that splits task-positive and task-negative regions would be more fully captured during movie-watching. Katsumi et al., (2021) have proposed FC gradients as a common neural architecture for predictive processing in the brain, with an emphasis on rest-G3 as a “model-precision gradient” (Katsumi et al., 2021). Considering that rest-G3 and movie-G4 appear to be based on the same underlying principle, the latter might provide a more refined or pronounced model for understanding these types of task-positive or predictive processes.

### 4.6. FC gradients across different movie stimuli are as reliable as across rest runs

Here, we demonstrate that the lower-dimensional gradient space is equally reliable for rest and movie despite the use of different movies during test and retest datasets (Fig. 4). This finding suggests that the variance captured by movie gradients reflects FC patterns characteristic of the naturalistic movie-watching experience on the whole rather than patterns evoked by a certain type of audiovisual stimulus. This may in part be due to the fact that each movie run was itself a compilation of different clips and FC values for each region capture the correlation of BOLD-signal time-courses across the full duration of the clips (i.e., they are collapsed, and not dynamic). Overall, this gradient finding follows suit with other work, including a meta-analysis across more diverse naturalistic paradigms, where a core set of common functional patterns are observed across even very different stimuli (Bottenhorn et al., 2019). This is not to say that there are no cross-movie differences in functional brain organization but rather that there are significant cross-movie similarities. The strong reliability of movie gradients across different movie stimuli could alternatively be interpreted as a low sensitivity of movie gradients to differences in the content/structure of different movies (e.g., intrinsic fluctuations could be obscuring movie-specific responses). Our findings do not address whether using the same movie across scans would result in higher test-retest reliability of gradient features. Additionally, it remains unclear whether different movies might reveal different gradients based on unique content, style, or idiosyncratic reactions to certain elements in a given movie, or whether regressing or filtering out non-stimulus driven activity might alter the gradients, their reliability, and/or the impact of different movies.

### 4.8. Naturalistic movie gradients provide more granularity for mapping cortical functional organization

Different types of data have been used to delineate organizational gradients in neuroscience, including structural, functional and evolutionary hierarchies (Sydnor et al., 2021). Direct neuroimaging measures such as inter-regional variation in the T1-weighted to T2-weighted ratios reveal a structural/anatomical hierarchy (Burt et al., 2018). Characterizing an evolutionary hierarchy in a neuroimaging-driven approach has quantified the ratio of cortical expansion between nonhuman primates and humans at each vertex (Krubitzer, 2007; Xu et al., 2020). Additionally, spatial gene expression measures have been shown to recapitulate FC during rest and to align with rest FC gradients (Huntenburg et al., 2018; Richiardi et al., 2015).

While gradient analysis of resting state or intrinsic FC measures has provided important insights into the functional organization of the cortex, resting state alone may not evoke a sufficient range of connectivity patterns to identify a functional hierarchy that accounts for diverse functional processes. As outlined in the Introduction, a long history of work suggests that brain organization follows a hierarchy from basic perceptual functions up to abstract cognitive processes. Studies that investigate dynamic and temporally embedded processes suggest that using ecologically valid data may be essential for delineating a hierarchy that takes into account complex, dynamic functions. For example, van der Meer et al., (2020) demonstrated that temporal brain state dynamics shift from predominantly bistable transitions between two relatively indistinct states at rest, toward a sequence of well-defined and highly reliable functional states during movie-watching that correspond with specific movie features (van der Meer et al., 2020). Another line of research also using naturalistic imaging paradigms has revealed a temporal integration hierarchy based on the time window during which presented information can influence the processing of incoming percept (Baldassano et al., 2017; Hasson et al., 2008). Notably, the topography of high inter-subject consistency necessary to describe temporal integration hierarchy aligns with the topography of high between-condition shift within gradient space (Baldassano et al., 2017) (Fig. 2B&D and Supplementary Video). Altogether, these observations suggest that movie-watching as a brain state may contain additional and unique information about the temporal dynamics and processing that are the result of evolutionary processes and demands. When used for gradient analyses, these data lead to more granular representation of the temporal and “in-action” functional hierarchy of the cerebral cortex.

In this Discussion, we have used descriptive nomenclature for the top movie gradients that is based simply on the cortical regions anchoring these gradients (i.e., sensorimotor, visual, auditory/language). These names summarize only the pole of the gradient, and there is some concern that the importance or utility of the “in-between” or liminal regions will be overlooked. In most cases, our findings suggest that the poles or extreme scores of a gradient will not be the most interesting or important. For example, the anchoring regions in the top movie gradients did not yield the strongest behavioral correlations (see Fig. 5 and Supplementary Fig. 6). Auditory regions did demonstrate some of the stronger correlations, but this was from within the middle range of the visual movie gradient. Similarly, visual regions demonstrated stronger brain-behavior correlations from within the middle range of the auditory/language movie gradient. Going forward, mid-gradient regions may be of particular interest when it comes to individual differences. We also recommend that descriptive names rather than gradient numbers be used for gradients when possible, as gradient order is likely to change across developmental and/or clinical groups.

The current study revealed important insights into the cortical organization under naturalistic conditions. However, our findings may have been limited by the exclusion of subcortical structures, especially given their integral role in coherent perception of the narrative of continuous stimuli such as movie clips (Baldassano et al., 2018; Cohen et al., 2022; Lee and Chen, 2022; Milivojevic et al., 2016). We also note that all of the movie clips in the HCP stimuli are relatively short in duration (less than 5 minutes each). This precludes capturing any temporal dynamics that may emerge over durations on par with a television episode (20-60 minutes) or feature film (1.5+ hours) which have been shown to be present and impactful on neural patterns that relate to memory and narrative constructions, etc. (Baldassano et al., 2017; Hasson et al., 2008; Zadbood et al., 2022). It would be interesting for future work to extend movie gradient investigations to longer movie scans with intact temporal and narrative continuity, or to directly test the effects of narrative continuity on gradient structure.

## 5. Conclusions

In the present study, we described cortical gradients of functional connectivity during movie-watching. The results show that during complex, dynamic processing, functional brain organization layers out into hierarchical gradients that are anchored by both default and frontoparietal regions, and that are unimodal. These naturalistic gradients thus follow different principles of cortical organization than are observed in resting state. In particular, because movie gradients are modality specific, they provide a more granular way to interrogate and understand principles of functional cortical organization. The progression of the sensory modalities captured by the top three movie gradients (by way of variance explained) follows known developmental and evolutionary hierarchies, from sensorimotor, to visual, to auditory/language regions. Future research could investigate developmental trajectories of these naturalistic gradients. We also show that movie gradient scores demonstrate stronger correlations with cognitive behavioral measures compared to resting state gradient scores. Movie gradients may offer unique advantages for future efforts to better understand cortical organization in neuropsychiatric disorders.

## Author contributions

**Ahmad Samara:** Conceptualization; Data curation; Formal analysis; Funding acquisition; Investigation; Methodology; Software; Validation; Visualization; Writing - original draft.

**Jeffrey Eilbott:** Data curation; Formal analysis; Methodology; Software; Writing - review & editing.

**Daniel S. Margulies:** Writing - review & editing.

**Ting Xu:** Methodology; Writing - review & editing.

**Tamara Vanderwal:** Conceptualization; Data curation; Funding acquisition; Investigation; Methodology; Project administration; Resources; Supervision; Writing - original draft; Writing - review & editing.

## Acknowledgments

We thank Boris Bernhardt and Pierre Bellec for helpful comments and discussion, and are grateful for the use of the Human Connectome Project 7T data and the BrainSpace toolbox. A.S. is supported by BC Children’s Hospital Research Institute (Brain, Behavior & Development Trainee Boost Award and Brain, Behavior & Development Doctoral Studentship). T.V. is supported by a BC Children’s Hospital Research Institute Investigator Award Program Clinician Scientist Award, and has funding from BrainCanada, SickKids Foundation and the Canada Foundation for Innovation John R. Evans Leaders Fund.

