## Supplemental tables and figures for "Cortical gradients during naturalistic processing are hierarchical and modality-specific"

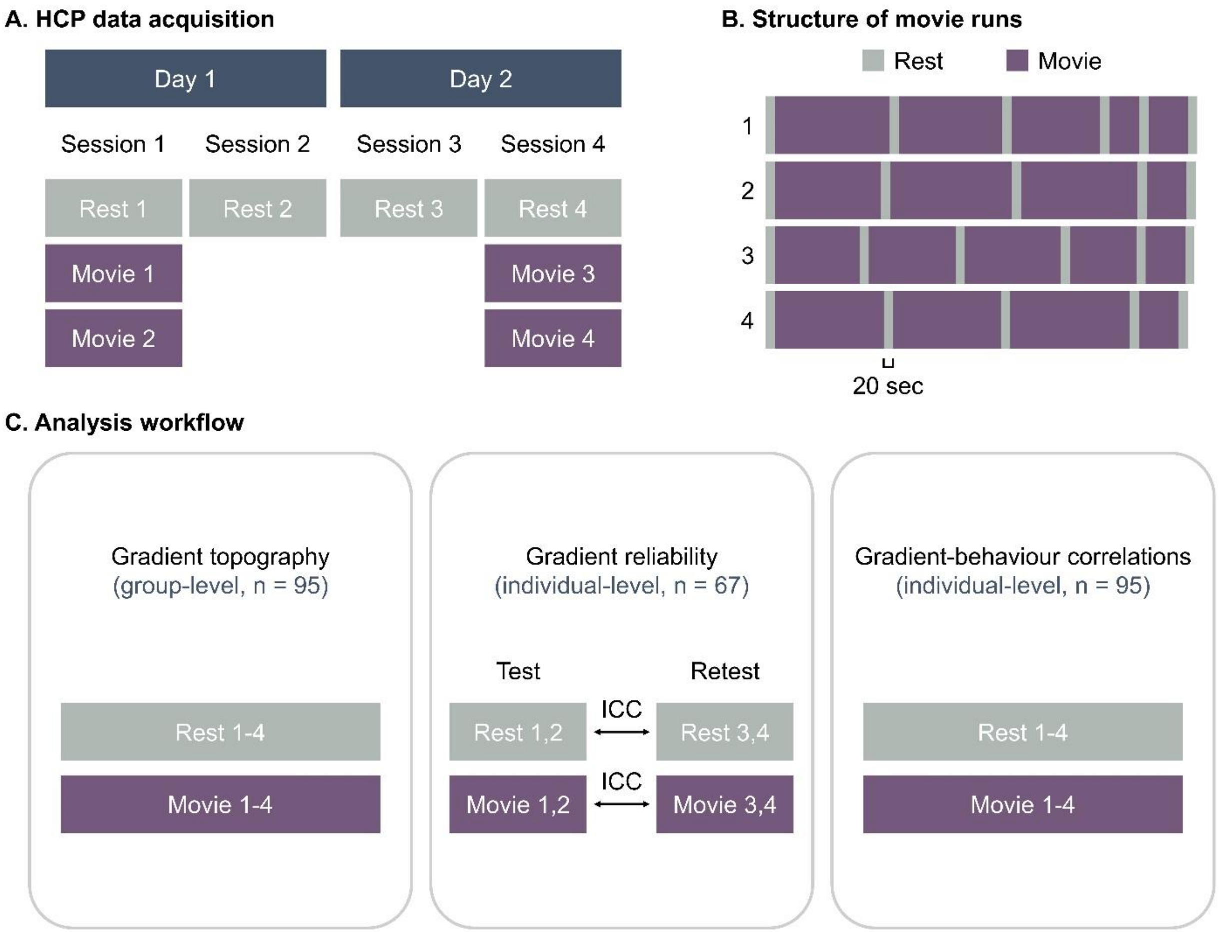


**Supplementary Fig. 1: Study design. (A)** Scan sessions from HCP 7T Release, showing how within-session and cross-day variance relates to each condition. **(B)** Within-run structure of movie-watching runs is shown, with both inter-clip and bookended rest epochs. **(C)** Workflow for each of the three major analyses is shown to highlight sample size differences and to show which data was used for which analysis.

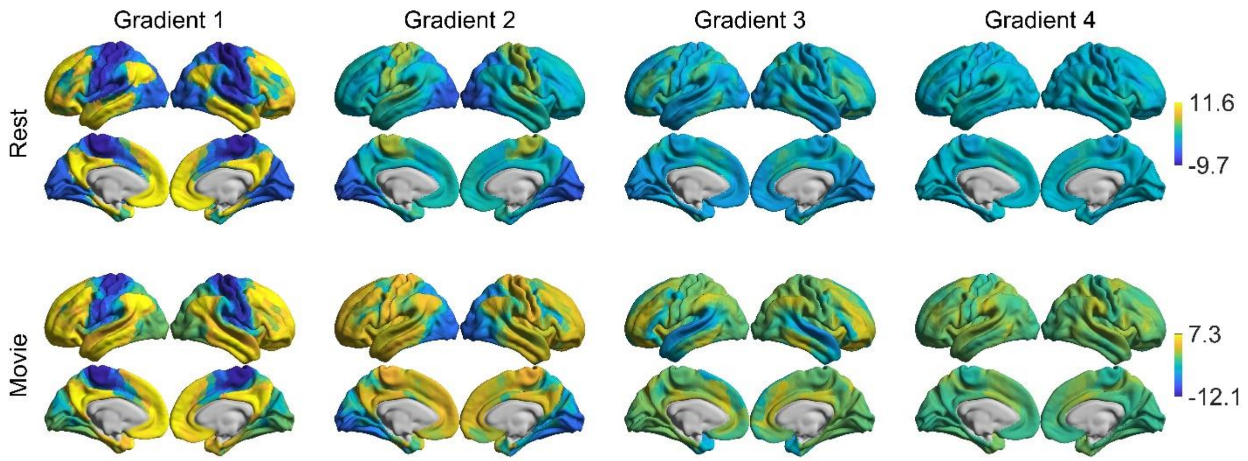


**Supplementary Fig. 2. FC gradients shown using the same scale within each condition (N=95, G1-4).** These are the same results shown in Fig. 1C where a different scale was used for each gradient. Here, the same scale is used to better represent the relative variance in Rest and Movie.

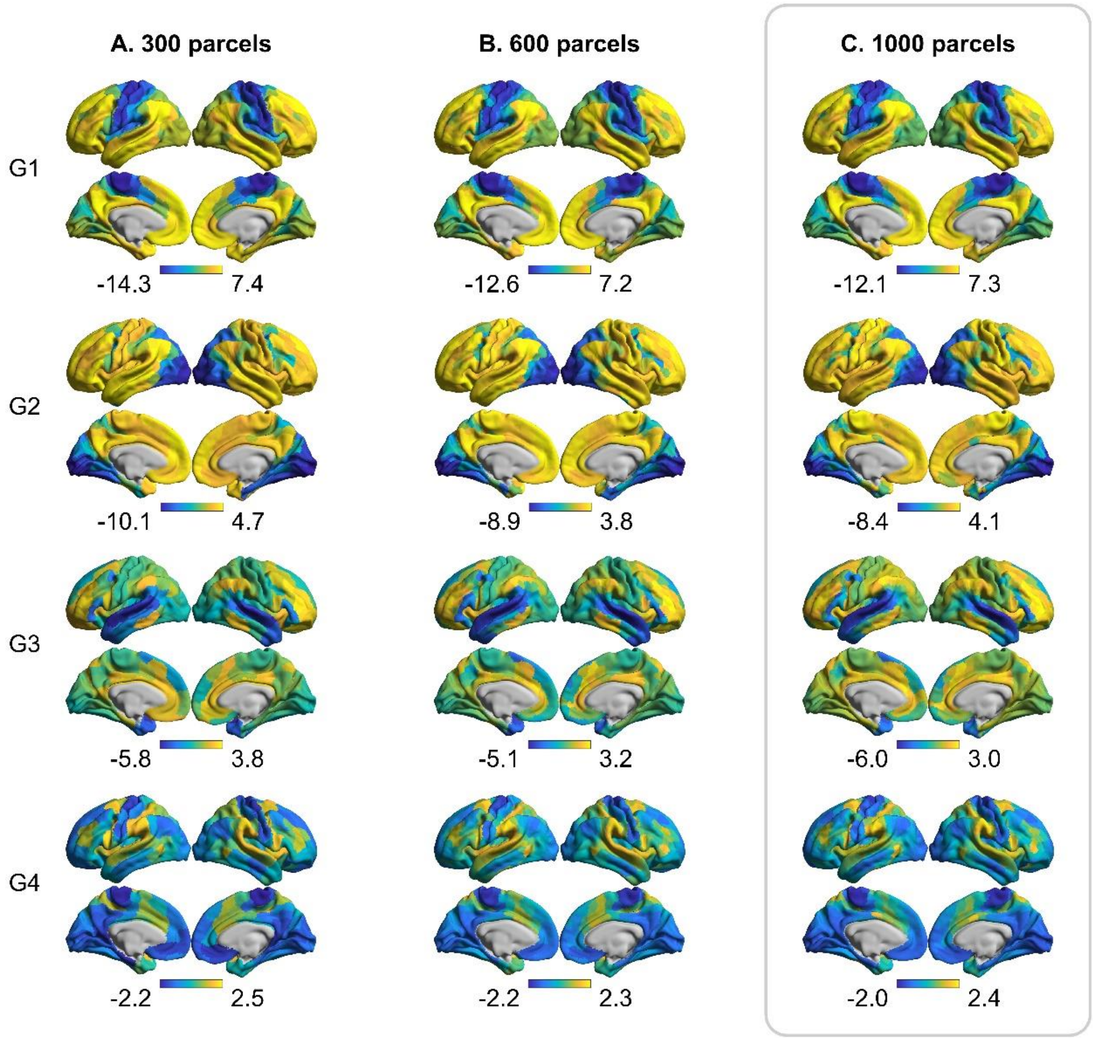


**Supplementary Fig. 3. Movie gradients across different parcellation resolutions.** These gradients were computed using the same gradient pipeline by varying only the spatial resolution from the Schaefer parcellation: 300 parcels **(A)**, 600 parcels **(B)**, and 1000 parcels **(C)**. The topography of movie gradients appears remarkably similar across the different resolutions. Schaefer 1000 (highlight in gray) was used in the main study.

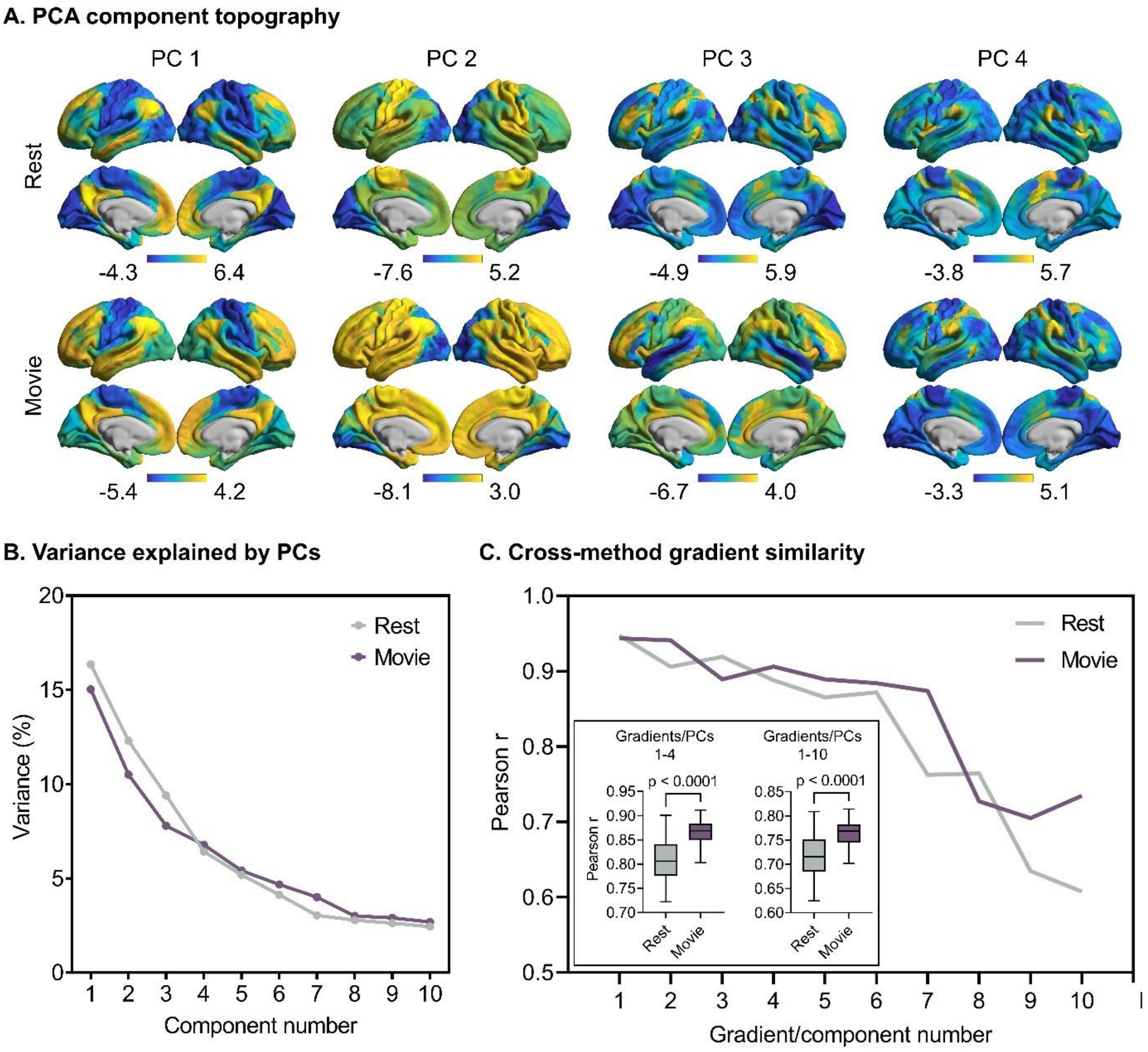


**Supplementary Fig. 4. PCA results (N=95, PCs1-4). (A)** Standard PCA was applied to thresholded (90%) group-averaged FC matrices for each condition. The topography of the component scores is highly similar to the diffusion embedding (DE) gradients (e.g., Fig. 1C). **(B)** Scree plot showing the variance explained by the first 10 PCA components. **(C)** Cross-method (PCA-DE) correlations are strong for each gradient within each condition suggesting that both linear and nonlinear reduction approaches recover the same lower dimensional organization. Inset shows t-tests for cross-condition differences for G1-4 and G1-10 of the cross-method correlations. Results from PCA and DE are more similar for Movie than for Rest.

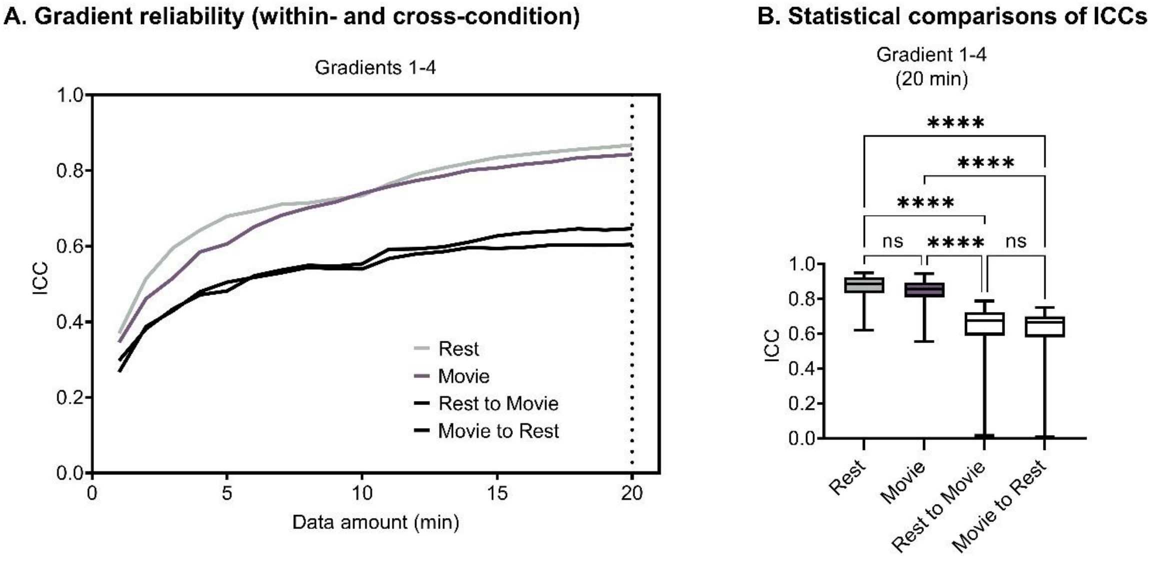


**Supplementary Fig. 5. Gradient reliability within- and cross-condition. (A)** ICCs for G1-4 when test and retest datasets are from the same condition (colored lines; also visualized in Fig. 4) and when the two datasets are from different conditions (black lines). **(B)** Within and cross-condition statistical comparisons of ICCs at 20 min of data. Within condition ICCs for both rest and movie were significantly higher than cross-condition ICCs in both directions, whereas no significant differences were observed between rest and movie or between the two directions of cross-condition reliability. ns = $P \geq 0.05$; **** = $P < 0.0001$.

**A. Topography of brain-behavior correlations (unthresholded)**

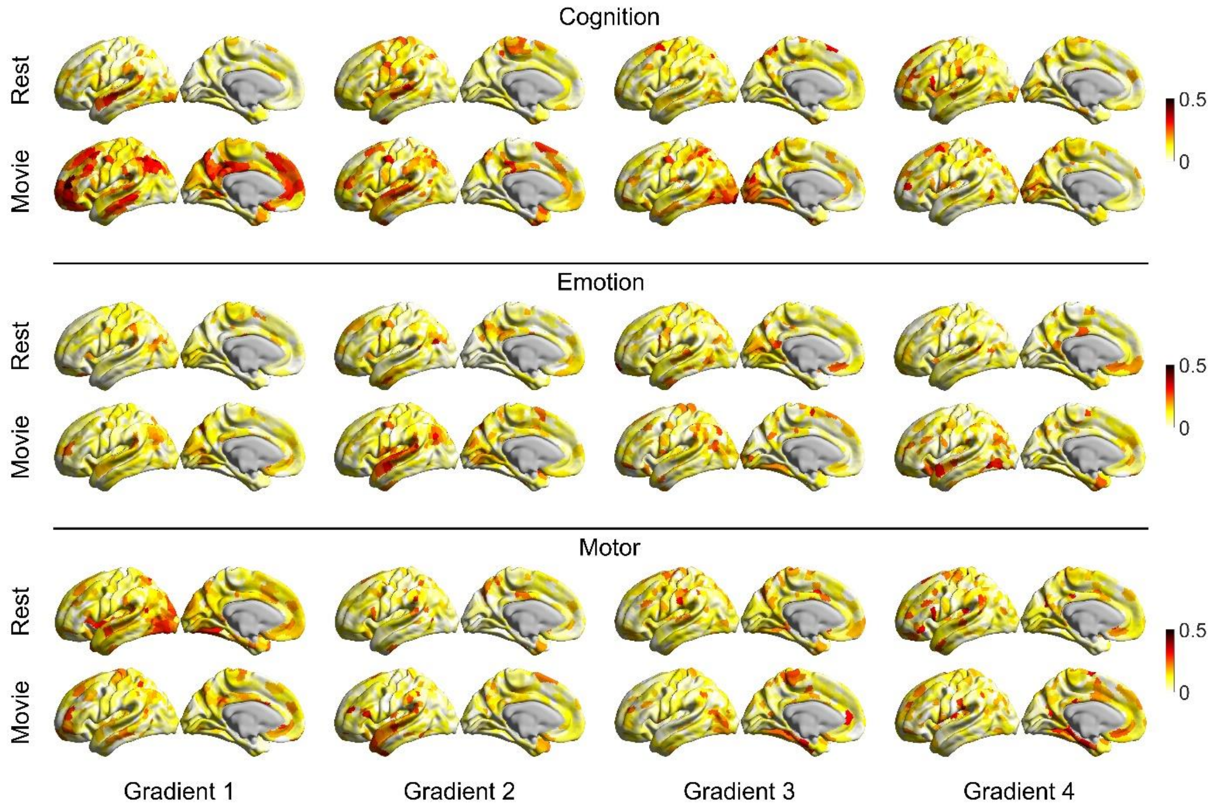


**B. Count of parcels |r| > 0.2**

| Rest | G1 | G2 | G3 | G4 |
|---|---|---|---|---|
| Cognition | 53 | 65 | 42 | 61 |
| Emotion | 34 | 36 | 87 | 30 |
| Motor | 162 | 55 | 89 | 85 |

| Movie | G1 | G2 | G3 | G4 |
|---|---|---|---|---|
| Cognition | 269 | 154 | 147 | 65 |
| Emotion | 55 | 117 | 72 | 89 |
| Motor | 69 | 56 | 99 | 91 |

**Supplementary Fig. 6. Unthresholded brain-behaviour correlations using gradient scores. (A)** Same data and procedure as shown in Fig. 5B, but here shown for all behavioural domains and unthresholded **(B)** Tables show count of regions with r > 0.2 for each gradient and condition.

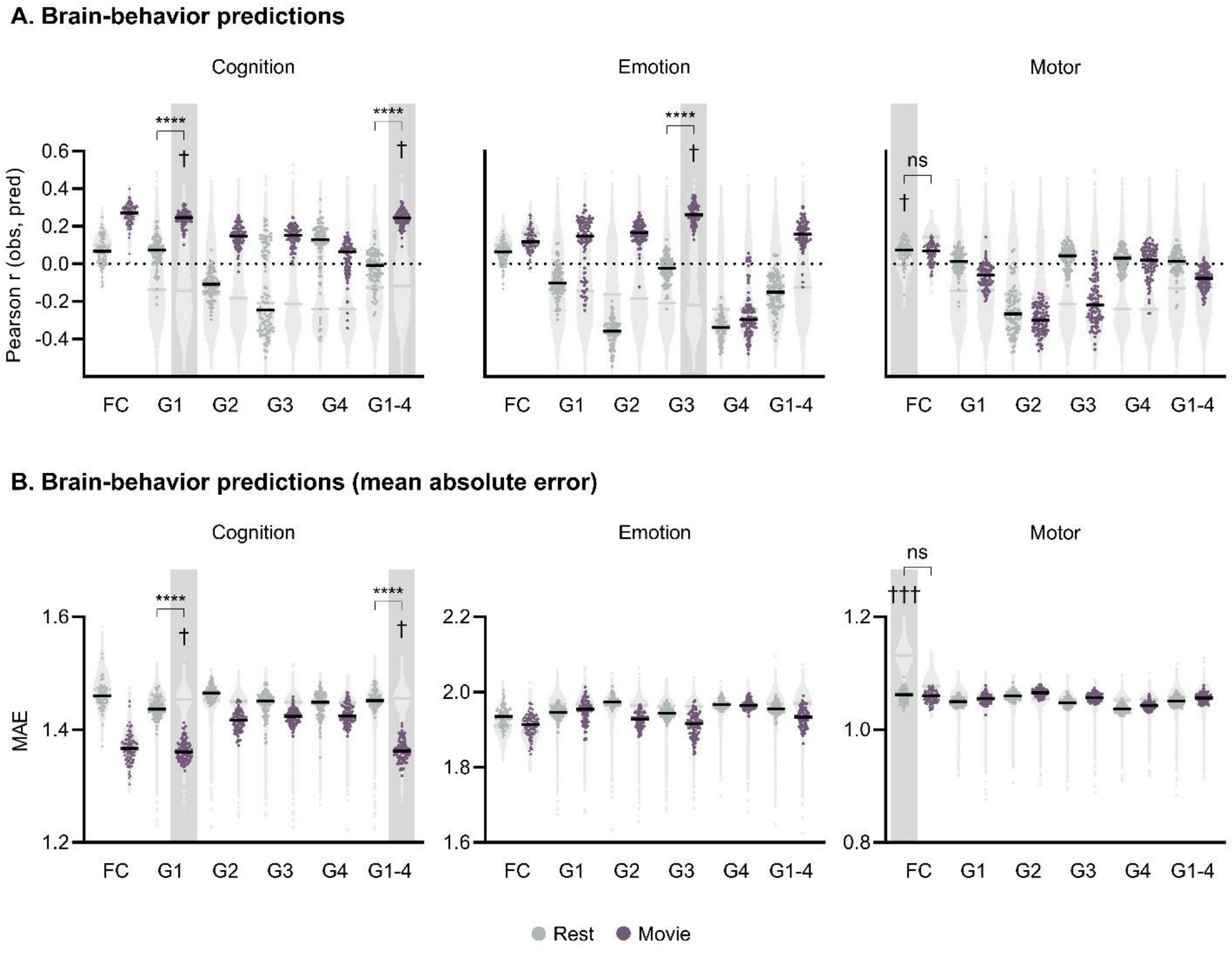


**Supplementary Fig. 7. Brain-behaviour predictions. (A)** Ridge regression was used for individual-level predictions of cognitive, emotion and motor scores. Model inputs are single gradient maps, combined gradient maps (G1-4), and full connectomes (i.e., FC matrices). Each dot represents the Pearson's r value between observed and predicted scores for one of 100 iterations. A null model (10,000 iterations) was generated for each model. Cross-condition differences in accuracy between Rest and Movie were assessed using t-tests for models that are significantly different from null models (highlighted in gray). **(B)** The same models are shown but using mean absolute error (MAE) as the accuracy measure. ns = P ≥ 0.05; **** = P < 0.0001; † = P < 0.05; ††† = P < 0.001.

**Supplementary Table 1.** PCA inputs for behavioural scores

| Measure | Short name | Domain |
|---|---|---|
| **Cognition** | | |
| Picture Sequence Memory | PicSeq_Unadj | Episodic memory |
| Dimensional Change Card Sort Test | CardSort_Unadj | Executive function/cognitive flexibility |
| Flanker Inhibitory Control and Attention Test | Flanker_Unadj | Executive function/inhibition |
| Penn Progressive Matrices | PMAT24_A_CR | Fluid intelligence |
| Oral Reading Recognition Test | ReadEng_Unadj | Language/reading decoding |
| Picture Vocabulary Test | PicVocab_Unadj | Language/vocabulary comprehension |
| Pattern Comparison Processing Speed Test | ProcSpeed_Unadj | Processing speed |
| Variable Short Penn Line Orientation | VSPLOT_TC | Spatial orientation |
| Short Penn Continuous Performance Test: Sensitivity | SCPT_SEN | Sustained attention |
| Short Penn Continuous Performance Test: Specificity | SCPT_SPEC | Sustained attention |
| Penn Word Memory Test | IWRD_TOT | Verbal episodic memory |
| List Sorting Working Memory Test | ListSort_Unadj | Working memory |
| **Emotion** | | |
| Penn Emotion Recognition: Number of Correct Responses | ER40_CR | Emotion Recognition |
| Penn Emotion Recognition: Correct Responses Median Response Time | ER40_CRT | Emotion Recognition |
| Anger-Affect Survey | AngAffect_Unadj | Negative affect |
| Anger-Hostility Survey | AngHostil_Unadj | Negative affect |
| Anger-Physical Aggression Survey | AngAggr_Unadj | Negative affect |
| Fear-Affect Survey | FearAffect_Unadj | Negative affect |
| Fear-Somatic Arousal Survey | FearSomat_Unadj | Negative affect |
| Sadness Survey | Sadness_Unadj | Negative affect |
| General Life Satisfaction Survey | LifeSatisf_Unadj | Psychological well-being |
| Meaning and Purpose Survey | MeanPurp_Unadj | Psychological well-being |
| Positive Affect Survey | PosAffect_Unadj | Psychological well-being |
| Friendship Survey | Friendship_Unadj | Social relationships |
| Loneliness Survey | Loneliness_Unadj | Social relationships |
| Perceived Hostility Survey | PercHostil_Unadj | Social relationships |
| Perceived Rejection Survey | PercReject_Unadj | Social relationships |
| Emotional Support Survey | EmotSupp_Unadj | Social relationships |
| Instrumental Support Survey | InstruSupp_Unadj | Social relationships |
| Perceived Stress Survey | PercStress_Unadj | Stress and self-efficacy |
| Self-Efficacy Survey | SelfEff_Unadj | Stress and self-efficacy |
| **Motor** | | |
| 2-minute walk | Endurance_Unadj | Endurance |
| 4-meter walk | GaitSpeed_Comp | Locomotion |
| 9-hole Pegboard | Dexterity_Unadj | Dexterity |
| Grip Strength Dynamometry | Strength_Unadj | Strength |

**Supplementary Table 2.** Full results of gradient-behaviour correlations

| Gradient | Rest mean r | Movie mean r | t | P |
|---|---|---|---|---|
| **Cognition** | | | | |
| Gradient 1 | 0.0727 | 0.149 | -20.78 | **< 0.0001** |
| Gradient 2 | 0.0917 | 0.1069 | -5.071 | **< 0.0001** |
| Gradient 3 | 0.0832 | 0.1127 | -9.706 | **< 0.0001** |
| Gradient 4 | 0.0851 | 0.0894 | -1.526 | 0.1273 |
| **Emotion** | | | | |
| Gradient 1 | 0.0774 | 0.0877 | -4.172 | **< 0.0001** |
| Gradient 2 | 0.0748 | 0.101 | -9.182 | **< 0.0001** |
| Gradient 3 | 0.0954 | 0.0907 | 1.524 | 0.1279 |
| Gradient 4 | 0.0735 | 0.0883 | -5.304 | **< 0.0001** |
| **Motor** | | | | |
| Gradient 1 | 0.1239 | 0.1001 | 8.182 | **< 0.0001** |
| Gradient 2 | 0.0766 | 0.0821 | -2.216 | **0.0269** |
| Gradient 3 | 0.1042 | 0.0959 | 2.824 | **0.0048** |
| Gradient 4 | 0.0966 | 0.098 | -0.4827 | 0.6294 |

**Supplementary Movie 1 hosted at the following link:**
https://raw.githubusercontent.com/tvanderwal/naturalistic_gradients_2022/main/resources/naturalistic_gradients.gif. Animation showing modulation of brain regions in gradient space from Rest to Movie. Below the scatterplot, cross-condition distance in gradient space is mapped onto the cortex to show that superior temporal sulcus regions move much further than other regions.
